# Disease resistance gene count increases with rainfall in *Silphium integrifolium*

**DOI:** 10.1101/2023.05.02.539110

**Authors:** Kyle Keepers, Kelsey Peterson, Andrew Raduski, Kathryn M. Turner, David Van Tassel, Kevin Smith, Alex Harkess, James D. Bever, Yaniv Brandvain

## Abstract

Intracellular plant defense against pathogens is mediated by a class of disease resistance genes known as NB-LRRs or NLRs (R genes). Many of the diseases these genes protect against are more prevalent in regions of higher rainfall, which provide better growth conditions for the pathogens. As such, we expect a higher selective pressure for the maintenance and proliferation of R genes in plants adapted to wetter conditions. In this study, we enriched libraries for R genes using RenSeq from baits primarily developed from the common sunflower (*Helianthus annuus*) reference genome. We sequenced the R gene libraries of *Silphium integrifolium* Michx, a perennial relative of sunflower, from 12 prairie remnants across a rainfall gradient in the Central Plains of the United States, with both Illumina short-read (n=99) and PacBio long-read (n=10) approaches. We found a positive relationship between the mean effective annual precipitation of a plant’s source prairie remnant and the number of R genes in its genome, consistent with intensity of plant pathogen coevolution increasing with precipitation. We show that RenSeq can be applied to the study of ecological hypotheses in non-model relatives of model organisms.

## INTRODUCTION

*Silphium integrifolium* Michx (commonly known as rosinweed or silflower) is a native perennial prairie species being domesticated for production as a forage and oilseed species (Vilela et al., 2018; Van Tassel et al., 2017). The plant’s perennial nature allows it to provide novel ecosystem services in agronomic production systems (Glover et al., 2010; Van Tassel et al., 2017) while producing oilseeds with similar nutritional composition to annual sunflower (*Helianthus annuus* L. (Asterales: Asteraceae) seeds (Reinert et al., 2018). However, *S. integrifolium* suffers infections from diverse pathogen enemies across its native range. The stems, leaves, and flowers can be infected with multiple strains (Turner, unpublished observations) of *Puccinia silphii* Schwein (1832), or *Silphium* rust (Turner et al., 2018), *Colletotrichum silphii* Davis (1919) (Horst 2008) syn. and *Colletotrichum dematium* (Pers.) Grove (Cybernome; Farr 1989) called leaf blotch hereafter. Recently, *Silphium* clear vein (SCV) leaf and stem disease has been described and hypothesized to be caused by a virus based on the identification of viral sequences in *Silphium integrifolium* that are similar to the Dahlia common Mosaic Virus (DCMV) and Dahlia endogenous plant pararetroviral sequence (DvEPRS, formerly DMV-D10) (Almeyda, 2014; Cassetta et al., 2023). Understanding how *S. integrifolium* defends against these pathogens is crucial for progress in breeding programs and will elucidate evolutionary questions posed in the species (Cassetta et al., 2023, Van Tassel et al, 2017). In this study, plants from 12 wild populations of *S. integrifolium* were planted in a reciprocal transplant design to better understand adaptation to disease pressures along a rainfall gradient. Phenotypic data revealed that these wild populations have distinctive morphologies and showed variable levels of tolerance to biotic stresses, such as pests and pathogens (Peterson et al., in prep). Individuals with resistance to diseases of *S. integrifolium* were visually identified in the study, and have been previously identified in breeding populations (Turner et al., 2018), but the genetic basis of disease resistance has not been investigated. Here, we look at the diversity of disease resistance genes in wild populations of *S. integrifolium* to answer evolutionary questions regarding adaptation of wild populations to their pathogen pressures and suggest applications for germplasm enhancement.

Environments with high precipitation are conducive to increased pathogen pressures (Clarkson et al., 2014; Granke and Hausbeck 2010; Islam and Toyota, 2004; Magarey et al., 2005; Rowlandson et al., 2015). Several studies have found patterns of resistance alleles consistent with pathogen selection intensity increasing with precipitation (Wahl 1970, Abbott, Brown, and Burdon 1991, Dong et al. 2009). We hypothesize that *S. integrifolium* populations evolved in areas with high precipitation will have a more diverse complement of disease resistance genes to deal with this challenge. To test this hypothesis, we collected seeds from *S. integrifolium* stands from four prairie remnants each in three geographic regions in the Central Plains of the United States. The regions are situated along a gradient of effective precipitation, or the amount of rainfall that is not lost to potential evapotranspiration, referred to in this study as the climate moisture index (CMI). We refer to these regions as “West”, from West-Central Kansas, “Central”, from Eastern Kansas, and “East”, from Central Illinois (Figure 1). Prairie remnants in the West receive around −55 kg/m^2^/month of CMI per year, whereas sites in the East receive closer to −15 kg/m^2^/month (Table 1). We note that plant defensive gene diversity increased along this gradient in *Andropogon gerardii*, a native perennial grass that co-occurs with *S. integrifolium* (Rouse et al. 2011) and soil pathogen diversity also increased along this same gradient (Delavaux et al. 2021).

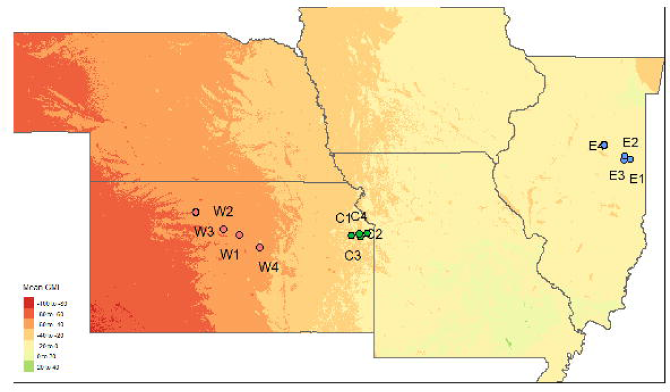

A major class of disease resistance genes in plants are the cytoplasmic nucleotide binding and leucine-rich repeat (NLR) genes. (Caplan et al., 2008; Eitas and Dangl, 2010). These NLRs, (hereafter, R genes), often contain an N-terminal domain that is either of a coiled-coil type, called CNLs, or a toll-interleukin-like type (TNLs), whose divergence predates the split between monocots and eudicots (Meyers et al., 1999), estimated to be about 160 Mya by TimeTree (Kumar et al., 2017). R genes are highly diverse, numbering in the hundreds in many plant species (Jupe et al., 2012; Toda et al., 2020). These genes evolve rapidly and are enriched in presence/absence variation even at the intraspecies level, limiting the utility of a single reference genome to elucidate the diversity of R genes within a species. This study employed R-gene enrichment sequencing (RenSeq; Jupe et al, 2013), which uses DNA baits to reduce complexity of genomic DNA libraries. The baits we designed come from the common sunflower, *Helianthus annuus* L., which is separated from *S. integrifolium* by between 22 (Meireles et al., 2020) and 33 million years ago (Zhang et al., 2021). We sequenced the RenSeq libraries from *S. integrifolium* plants across the rainfall gradient using both Illumina short-read and PacBio long-read technologies.

RenSeq is not a new technology – having been developed in 2013. It has been used to address basic problems in plant immunity (Jupe et al., 2013) and applied questions in plant breeding in numerous crop species and their wild relatives for crop improvement (e.g. Arora et al., 2019) and to understand mechanisms of plant immunity (e.g. Witek et al., 2021). Despite the utility of RenSeq in plant breeding projects, and in helping us better understand plant disease resistance, this technology has not been used to address basic evolutionary and biogeographical questions.

Using both simple linear models as well as mixed effect models, we found a significant positive correlation between the effective precipitation of a plant’s host prairie and the number of R genes detected in both the Illumina and PacBio datasets. While this observation does not itself prove the broader theory that disease resistance correlates with rainfall in plants, it is certainly consistent with theoretical expectations, and should motivate additional evaluation of these ideas in other taxa. Because we demonstrate the economic utility of the RenSeq approach in non-model systems, our work not only points to data consistent with our hypothesis but paves the way towards evaluating the generality of this finding.

## METHODS

### Plant Materials

*Silphium integrifolium* seeds were collected from 12 prairies in 2017 and 2018 (Figure 1; Table 1). The prairie remnants collected spanned from central Kansas to central Illinois. The prairie remnants have variable historical, geographic, and climatic characteristics described in Cassetta et al 2023. From each of the 12 prairies, seeds were collected from approximately 40 mother plants randomly chosen from throughout the prairie remnant. As *S. integrifolium* is an obligate outcrosser, seeds collected from a single mother were a half sibling family.

*S. integrifolium* seeds were germinated at room temperature together in a heated greenhouse at The Land Institute, Salina KS in 2019. A total of 96 genets comprising eight individuals from each prairie remnant from different mothers were selected from the seedlings. Selection of the 96 individuals was randomized although the seedlings chosen had to be moderately vigorous. The pots containing the seedlings were randomized and placed on greenhouse benches. Pots were re-randomized several times during the season. The plants grew rapidly but remained as vegetative rosettes. Each crown (genet) was divided into 4 pieces (ramets) with approximately equal numbers of dormant buds and roots, although a few crowns were too small and only produced 2-3 pieces. The ramets were repotted into 10-gallon fabric planters (Smart Pot) using Pro-Mix BX.

### Tissue Collection

One ramet from each genet was grown in The Land Institute Greenhouse until young leaves had emerged. On March 12, 2020, black plastic pots were placed over one clone of each of the 99 plants emerging rosette and left for 48 hours to slightly etiolate the young leaves, which improves the quality of extracted DNA in photosynthetic active tissue containing phenolic compounds (Dabe et al., 1993). On March 14, 3-5 unexpanded leaves were excised from each crown. The collected tissue was placed in a labeled envelope and immediately placed on dry ice in a cooler. Within 60 minutes of collection, the sampled leaf tissue was placed in a lyophilizer and lyophilized for 48 hours.

### Clone placement across the gradient and phenotyping

After young leaf tissue was collected from plants in the greenhouse, the plants remained in the greenhouse under ideal conditions allowing them to grow rapidly. One cloned ramet from each of the available genets was then moved to each of the following locations within the regions the seed was sourced from: 1. breeding plots in Salina, KS at The Land Institute (West), 2. Prairie Park Nature Center, Lawrence, KS (Central), 3. research plots at the ISU horticulture center in Normal, Illinois (East), and 4. Land Institute Crop Protection Ecology greenhouse (backup site to conserve genets in a disease and insect free state). The pots remained undisturbed at each site for two years, 2020 and 2021 except the Central site cloned plants, which, in May 2021, were moved from the Prairie Park Nature Center to the Perennial Agriculture Project field station, approximately, 6 miles away. The clones received occasional watering in each site to promote disease growth and avoid plant death due to the above ground pots drying out.

In 2020 and 2021, disease scores and insect herbivory damage were recorded on each of the clones in all 3 sites. Data was collected during their flowering period, between the months of July and September. Diseases were recorded according to methods in Cassetta et al., 2023. Briefly, leaf rust severity was quantified by visually assessing the total percentage of leaf area affected (0-100%) using the rust scale described by Turner, et al. (2018). Similarly, stem rust severity was recorded as the total percentage of the stems covered in rust (0-100%). Leaf blotch was quantified by assessing the total percentage of leaf area affected (0-100%). Silphium clear vein was rated on a 0-10 scale based on symptoms observed on the entire plant including: leaf distortion (narrow width, curled, toothed, twisted, or pinched tips), and twisted meristem symptoms. Leaf herbivory scores were collected by quantifying the percentage of total leaves with evidence of insect herbivore (mining, sucking, chewing) damage (0-100%).

### Climate Moisture Index

A BIOCLIM+ raster map of the continental U.S.A. containing 30-year climate moisture index averages (1981-2010) was downloaded from the CHELSA (Climatologies at high resolution for the Earth’s land surface areas) database (Brun et al., 2022). The climate moisture index is calculated as the difference between mean annual precipitation and mean potential evapotranspiration and, when negative, represents the availability of water as a limiting resource in dry regions such as prairies.

### Sequencing/RenSeq/QC

DNA was extracted from lyophilized leaf tissue using the CTAB method by the University of Minnesota Genomics Center. DNA was then shipped on dry ice to Arbor Biosciences for sequencing, library preparation, and RenSeq data acquisition. Extracted DNA was prepared for sequencing using Illumina library prep kits. Sequencing libraries were enriched for R genes using the RenSeq protocol described in Jupe et al., 2013 using Arbor’s myBaits systems. In short, biotinylated single-stranded RNA oligo baits were designed based on annotated R genes from *Helianthus annuus*, a relative of *Silphium integrifolium*, within the same subtribe, estimated to be between 22.5 (Meireles et al., 2020) and 33.5 million years divergent (Zhang et al., 2022). Hybridized oligos were pulled down using streptavidin-coated magnetic beads. Libraries were then sequenced on an Illumina NovaSeq S4 sequencer using paired-end 150bp reads by Arbor Biotechnologies in Cambridge, MA.

A subsample of 10 individuals was selected for a complementary R gene analysis using Pacific Biosciences (PacBio) long-read sequencing. High-molecular-weight enriched DNA from three individuals from each of the West, Central, and East prairies, as well as one individual derived from a cultivated lineage, were prepared for long read sequencing and then sequenced on a Pacific Biosciences Sequel II using circular consensus sequencing (CCS) by Arbor Biotechnologies.

A total of 99 Illumina libraries (96 from prairie remnants + 3 cultivated “elite” lineages) and 10 PacBio libraries were sequenced. Both library types were filtered for sequence quality using Trimmomatic v0.39 (Bolger et al., 2014) and then assessed using fastqc v0.11.7 (Andrews 2010). The Illumina libraries were subsampled to a standardized assembly input size of 1Gb to avoid biasing R gene counts by library input size. The PacBio libraries were not subsampled to a standard input size, but a correlation between input size and total R gene count in this data type reveals no dependence of final assembly size on initial input size (Figure S3; p=0.414). Six Illumina libraries failed to meet the input threshold and were excluded from analysis. The libraries that met the input threshold were then *de novo* assembled using SPAdes v3.15.3 (Bankevich et al., 2012) with -k 21,33,45,55,65,81.

### Draft Genome Assembly

A chromosome-scale genomic assembly of an F1 *Silphium integrifolium* x *S. perfoliatum* is being prepared at the time of writing. The *S. integrifolium* mother was a leaf blotch resistant, large-headed selection from a breeding population that had undergone approximately 6 cycles of recurrent selection for increased number of fertile ray florets (“feminization”) per head and disease resistance. The founders of the breeding population were sourced from wild central Kansas populations, i.e., located within the geographic region we call “West” in this study. Flash-frozen tissue was sent to Arizona Genomics Institute for high-molecular-weight DNA extraction, which was then prepared for PacBio Sequel-II CCS sequencing at the Genome Sequencing Center at HudsonAlpha Institute for Biotechnology. The libraries were sequenced at roughly 21.5X depth assuming a 10 Gb genome size. Reads were assembled *de novo* using hifiasm assembler (Cheng et al., 2021) using default parameters.

### R Gene Annotations

We annotated the Illumina RenSeq libraries, PacBio RenSeq libraries, and the draft genome for their R gene content. Each assembly was annotated for R genes using NLR-Annotator (Steuernagel et al., 2020), which searches for common R gene peptide motifs in protein-coding sequence. The annotations were performed with the “-a” flag, which outputs the amino acid sequence of the nucleotide-binding domain (NBARC domain). We used the NBARC sequences of the draft genomic assembly to infer a maximum likelihood tree in RAxML v.8.2.11 (Stamatakis 2006; Stamatakis 2014) using the peptide substitution model “PROTGAMMAAUTO” with 100 rapid bootstraps.

The two broad, non-overlapping classes of R genes that we analyzed in this study, TNLs (containing a toll-interleukin-like type N-terminal domain) and CNLs (containing a coiled-coil type N-terminal domain), were identified from the annotations based on diagnostic motifs. To identify TNLs, we parsed annotations for the motifs 13, 15, and 18 (as defined in Table S1 of Jupe et al., 2013). For CNLs, we searched for motifs 2, 6, 16 and 17. These two sets of motifs occur only in their respective class of N-terminal domain.

### Principal Coordinates Analysis

To test for associations between R genes and the phenotypes collected on the cloned plants across the gradient, we performed simple linear modeling as well as mixed effect modeling, which treats CMI as a fixed effect, and source prairie as a random effect. We found one significant correlation between R Genes of the various populations and leaf blotch. To interrogate which R genes were responsible for the association between leaf blotch and R gene counts, we performed a census of which R gene NBARC sequences occurred in which individuals (Table S5). The census was then used to generate a principal coordinate analysis in the R package “vegan” (Oksanen 2015) against each of the five phenotypic covariates included in the study. P-values were corrected for multiple tests using the Benjamini & Hochberg method (1995).

### Library Enrichment Calculations

The fasta files containing the annotated R genes for each Illumina library were concatenated into a fasta containing every observed R gene from the Illumina dataset (hence, a “Pan-NLRome”). Reads from each Illumina RenSeq library, as well as eleven 1X-depth WGS libraries from seven *S. integrifolium*, two *S. perfoliatum*, one *S. laciniatum* and one *S. terebinthinaceum*, were mapped to the Pan-NLRome using *bwa mem* (Li 2013). The fraction out of the total reads in the library that mapped to the Pan-NLRome with a SAM CIGAR mapping score of between 148M-151M was calculated, representing reads that mapped to an R gene in the Pan-NLRome with between 98.01%-100% identity.

## RESULTS

### Reference Genome Assembly

The draft genome used in this study was 9.9 Gb in length and totalled 2,584 contigs. The longest contig was 182 Mb long and the mean contig size was 3.8 Mb. The L50 (which is the number of contigs, ordered longest-to-smallest, to comprise 50% of the assembly size) was 77, and the N50 (the size of the contig that passes the 50% assembly size threshold) was 40.2 Mb.

The maximum likelihood tree constructed from the R genes in the draft assembly was annotated according to peptide motifs that are diagnostic for either CNL or TNL identity (Figure 2). A deep evolutionary divergence between the two broad classes is seen, although we observe isolated instances of R genes falling incongruently within the wrong clade.

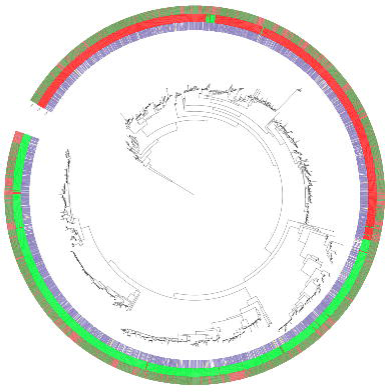

### Library Enrichment

The libraries of seven *S. integrifolium* that were sequenced using WGS, and thus were unenriched for R genes, contained a median of 1.78% of reads that mapped with high fidelity (98% identity or higher) to contigs in the Pan-NLRome. Libraries that had been enriched for R genes using oligo baits designed from *Helianthus annuus* contained a median of 63.4% of high fidelity reads mapping to R genes, representing a 36-fold enrichment for R genes (Figure 3).

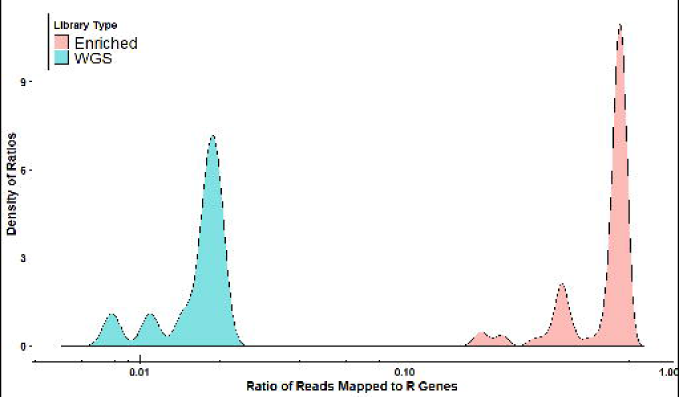

### R Gene Counts by Geographic Region

The Illumina R gene assemblies ranged from 2.2 Mb to 51 Mb, and averaged 15 Mb in size. We detected an average of 386.3, 410.0, and 414.0 R genes in plants sourced from the Western, Central, and Eastern prairies, respectively (left facet of Figure 4A). The plants averaged 47.38 (West), 48.69 (Central), and 45.14 (East) TNL genes, and 166.1 (West), 171.3 (Central), and 175.3 (East) CNL genes (Table S2).

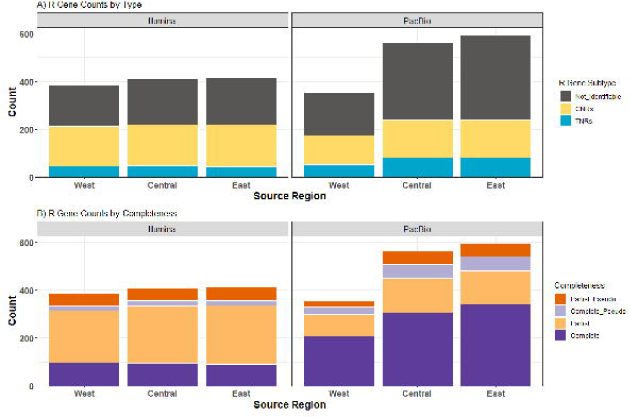

The PacBio R gene assemblies ranged from 4.9 Mb to 13 Mb, but averaged 8 Mb. Within this dataset, we only had three plants per region, so the inferential power was lower, but our confidence in the contigs assembled was higher due to longer reads. We found an average of 355.0, 563.3, and 592.7 R genes in the Western, Central, and Eastern prairies, respectively (right facet of Figure 4A). Within this dataset, we found an average of 54.00 (West), 81.67 (Central), and 81.33 (East) TNLs, and 121.3 (West), 158.7 (Central) and 159 (East) CNLs (Table S2). As compared to the Illumina data, PacBio assemblies resulted in more “complete”, as compared to “partial” genes for both intact and pseudogenized R genes (Figure 4B), indicating that many of the R genes predicted as pseudogenes in the Illumina data likely represent complete genes that are mis-categorized due to assembly limitations.

### Comparing to Reference Genome

The reference genome, which was sequenced at 21.5 X coverage using PacBio long reads, contained a greater number of R genes in total (873; Table S2), of which 281 and 147 where annotated as CNLs and TNLs, respectively. This compares to the 603 candidates found by NLR-Annotator in the closely related sunflower genome (Toda et al., 2020). It contained a comparable percentage of complete R genes relative to its total number of R genes (50.7%) as the enriched PacBio libraries (58.2%, 54.2%, and 57.3% in the West, Central, and East, respectively). 837 R genes in the reference (as well as the 281 and 147 CNLs and TNLs, respectively) exceeded counts from both the lower-coverage, R-gene-enriched PacBio assemblies or the Illumina short-read assemblies, below, suggesting either that the reference had more R genes (which is plausible as it an interspecific F1) than the rest of the samples and/or that some genes were missed in the enrichment process.

### R Gene Counts Increase with Increasing Climate Moisture Index

We found a strong positive association between R gene count and CMI in both the Illumina and PacBio datasets (Figure 5A,B; Table S3) from the simple linear model. We also found a significant positive correlation between Illumina R gene counts and CMI using the mixed effect model (MS_CMI_ = 9493, MS_error_ = 1059.6, F = 8.96. Degrees of freedom from Satterthwaite’s method [Satterthwaite 1946]: df_CMI_ = 1, df_error_ = 9.73, p = 0.0139, Figure 5A; Table S4), on top of substantial variation among source prairies. Specifically, the model predicted an additional 0.707 kg/m^2^/month (with a standard error of 0.236 kg/m^2^/month) R genes for every additional unit of CMI. As average CMI ranged from −57.3 units to −12.9 units, the model predicts that plants from prairies with the highest annual CMI will have 31.4 more R genes than those from prairies with the least CMI. Because most variation in CMI is associated with source region (West, Central, or East), we also modeled source region, rather than CMI as a fixed effect. This analysis revealed a similar, but nominally insignificant association between source region and the number of R genes (MS_region_ = 8753, MS_error_ = 4376.5, F = 4.13. Degrees of freedom from Satterthwaite’s method: df_region_ = 2, df_error_ = 8.82, p = 0.0542, see descriptive statistics in the section above and in Table S4).

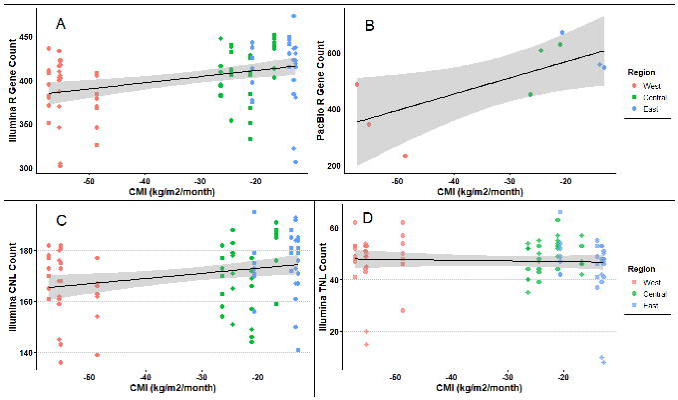

Our PacBio sequencing recovered more R genes per individual and had relatively fewer R genes annotated as “partial” (relative to “complete”) than the Illumina data (Table S2). Despite much lower power, we still see a significant association between CMI and R gene count in the PacBio data set (MS_CMI_ = 92006, MS_error_ = 9961, F = 7.193, df_CMI_ = 1, df_error_ = 7, p = 0.0315, Figure 5B). Like the case for the Illumina data modeling R gene count from the PacBio data revealed a similarly consistent but nominally insignificant association (MS_region_ = 53074 MS_error_ = 10164, F = 4.96. Degrees of freedom df_region_ = 2, df_error_ = 6, p = 0.054, see descriptive statistics in the section above and in Table S4).

### CNLs and TNLs

We did not find a significant relationship between CMI and TNL counts (p = 0.695, Figure 5D, MS_region_ = 14.072, MS_error_ = 86.559, F = 0.1626. Degrees of freedom from Satterthwaite’s method: df_region_ = 1, df_error_ = 10.215), and only a marginal significance with CNL counts (Figure 5C; p=0.0634, MS_region_ = 651.19, MS_error_ = 150.22, F = 4.3349. Degrees of freedom from Satterthwaite’s method: df_region_ = 1, df_error_ = 10.22).

Due to having only one plant sampled per prairie, we report the results of a fixed-effect linear regression model for the PacBio dataset. Despite a much lower inferential power (n=9), we still found a positive correlation between CMI and the PacBio R genes (Figure 5B; p=0.019). There was a significant relationship between CMI and PacBio TNL counts (Figure S1B; p=0.018), but not CNLs (Figure S1A; p=0.14).

### Correlations Between Phenotypes and R Gene Counts

There were no significant correlations between Illumina R gene counts and leaf rust severity (Figure 6B; p=0.63), rust response (Figure 6C; p=0.73), stem rust severity (Figure 6A; p=0.51), or insect leaf herbivory (not pictured; p=0.23). We found that R gene counts were significantly associated with leaf blotch scores (Figure 6D; p=0.0043). When source prairie was modeled as a random effect, leaf blotch scores were still found to significantly correlate with CMI (p=8.94e-5, slope=-0.0082).

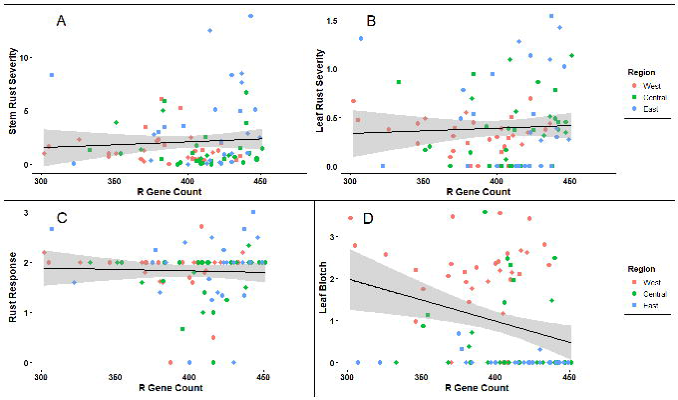

### Principal Coordinate Correlations Among Phenotypes

The PCoA analysis (Figure S4) of the unique NBARC sequences (Supplemental file “AllNBARCs.fasta”; Table S5) against the disease phenotypes yielded no significant candidates for genes responsible for disease resistance after correcting for multiple tests.

## DISCUSSION

We enriched *Silphium integrifolium* libraries for R genes using baits developed from the related *Helianthus annuus* genome to characterize the diversity across a rainfall gradient. We found that the baits recovered a substantial portion of R genes in the reference. We found geographic associations between R gene count and rainfall. Here, we discuss the *Silphium* R gene content across the rainfall gradient, as well as the performance of the RenSeq enrichment method, and the biological implications of the geographic pattern we uncovered.

### DISEASE RESISTANCE AND RAINFALL

#### R Gene Counts Correlate with Precipitation

The significant correlation between net precipitation rate, as estimated by average CMI, and the number of R genes in a plant’s genome, detected in both the Illumina short-read dataset as well as the underpowered PacBio long-read libraries (n=9), suggests evidence of possible genomic adaptation for higher pathogen load. In fact, the steeper relationship of R gene number and CMI observed with the PacBio long-read libraries suggests that the Illumina dataset underestimates the strength of this relationship. The trend does not appear to be driven by either TNLs or CNLs, for which the results conflict between the Illumina and PacBio datasets. Genes of type TNL, or those containing a toll-interleukin-like receptor in their N-terminal domain, don’t differ significantly according to CMI. A stronger understanding of the relationship between climate factors and the evolution of these two subtypes of R genes remains to be investigated.

#### Mixed Disease Phenotype Results

Despite the significant trend of the number of R genes increasing with CMI, leaf rust severity, leaf rust response, and stem rust severity did not significantly vary in the common gardens. These three traits are different phenotypic metrics to assess a specialist fungal pathogen, *Puccinia silphii*, commonly known as *Silphium* rust. These results are consistent with results from common gardens that found that eastern populations of *S. integrifolium* were not, on average, more resistant to *Silphium* rust (Cassetta et al., 2023). One possible explanation for why resistance to *Silphium* rust was not associated with variable R gene counts may lie in the concept of the ‘disease triangle’ reviewed by Velásquez et al., 2019. The clones in this study were placed in environments where *S. integrifolium* pathogens were present, and experienced conditions that would be ideal for pathogenic growth: warm and wet. Even with effective immunity represented by the R genes sequenced in the plant populations, the level of resistance conferred by the resistance genes might have been obscured by the pathogen spread. Disease resistance in these wild populations is highly likely to be quantitative, meaning that resistance is not an all-or-nothing response, but rather a distribution of potential responses ranging from minor to major (French et al., 2016). Further studies of the wild populations with sequenced R genes, with controlled inoculations of plants with pathogens, would elucidate associations between R genes that may confer various levels of response to pathogens.

Further, the evolutionary dynamics of R genes may be muddled by seemingly competing forces of selection, with “arms race” dynamics expected to generate gradients in total host resistance, while oscillatory “red-queen” dynamics may not (Bergelson et al., 2002). It is possible, for example, that local co-evolution of *S. integrifolium* and *Silphium* rust has resulted in adaptation of the rust to its local host population, rather than variation in overall resistance. This red-queen dynamic is supported by observations from common gardens distributed along the gradient that showed much greater virulence of *Silphium* rust on their populations from their local region than from other regions (Cassetta et al. 2023). In the case of *Silphium* rust, R gene identity may be more important than R gene diversity.

We did observe that resistance allele diversity was associated with suppression of leaf blotch (Fig. 6d). Our result affirms observations of a previous common garden study that found that eastern populations of *S. integrifolium* were more resistant to *Silphium* blotch, as well as to clear vein virus (Cassetta et al. 2023). These results are consistent with expectations of greater diversity of resistance alleles contributing to greater resistance to pathogens, and are qualitatively consistent with arms-race-type dynamics. In *S. integrifolium*, leaf blotch is caused by the generalist fungal pathogen *Colletotrichum dematium* (Cybernome; Farr 1989), as well as its more specialized congener, *Colletotrichum silphii* Davis (Horst 2008). It is possible that arms race dynamics are more likely between hosts and generalist pathogens such as leaf blotch, while specialist pathogens such as the Silphium rust are more likely to generate red-queen dynamics.

We note that the observed relationship between resistance allele diversity and resistance to blotch may be spurious, as other heritable factors that covary with rainfall might contribute to disease resistance, such as plant secondary chemicals. We note, however, that insect herbivory may be an indicator of such overall resistance, and we did not observe any relationship between R gene diversity and insect leaf herbivory. This result is consistent with R genes predominantly acting as pathogen effector receptors, and not targeted toward resistance for insects (Chovelon et al., 2021).

### A BROADER PATTERN OF R GENE EVOLUTION

#### R Gene Genealogy Recovers Expected Patterns

The maximum likelihood genealogy constructed from the NBARC domains of the R genes of the draft genomic assembly recovers the deep evolutionary divergence between CNL-type and TNL-type R genes that has been reported in many other studies (Meyers et al., 2003; Mun et al., 2017; Neupane et al., 2018). The instances of individual R genes nesting incongruently within the wrong clade may be attributable to N-domain switching via recombination, which, to the best of our reckoning, has yet to be observed. More likely, it may be due to misidentification of upstream nucleotide sequence as incidentally translating to peptide sequence that meets the identity threshold for one of the diagnostic motifs that places those R genes in the wrong categorization.

### PERFORMANCE OF RENSEQ FOR R GENE ENRICHMENT

#### Enrichment Using Distant Reference

The R gene libraries presented in this study were developed by enriching *Silphium integrifolium* DNA for R genes using baits developed from *Helianthus annuus*, a model organism with well-developed genomic resources. Despite an estimated divergence age of between between 22.5 (Meireles et al., 2020) and 33.5 million years ago (Zhang et al., 2022), the baits were able to enrich libraries to contain a median of 63% of reads originating from R genes, representing a 36-fold enrichment over WGS. For comparison, a previous study by Andolfo et al. (2014) found success enriching the *Solanum lycopersicum* (common tomato) with baits designed from *Solanum tuberosum* (common potato), a congener estimated to be 6.7 Ma divergent by TimeTree (Kumar et al., 2017). This study demonstrates the economical promise of RenSeq for studying the immune systems of non-model plants under a variety of ecological and evolutionary pressures. It also showcases the screening of crop wild relatives that are of agronomic interest, such as *S. integrifolium*, for disease resistance genes that might enable more robust response to pathogens that pose a challenge for the domestication of the plant.

The reference genome contained a much higher number of R genes in the draft genome assembly compared to the enriched libraries (873 compared to ∼400-600), which can be accounted for by two factors. First, the draft genome is assembled from an F1 hybrid between two different species, *S. integrifolium* and *S. perfoliatum*. While we expect overlap in many of the R genes due to homology, our count is likely an overestimate of the true number contained within the haploid genome of *S. integrifolium* due to the nature of R genes as a rapidly diversifying gene family. Second, RenSeq enrichment likely captures a different (and smaller) subset of the R genes than the WGS PacBio sequencing we employed for the draft assembly.

#### Caveats and Future Directions

Consistent with the prediction that more rainfall results in more pathogens, which result in more disease resistance – we find a positive correlation between R gene counts and precipitation. However, the distance between our observed correlation and our causal motivation is quite large. Here we suggest two routes to bridge this gap – independent replication and unraveling the causal chain.

Correlation does not imply causation. However, repeated independent replication “does waggle its eyebrows suggestively and gesture furtively while mouthing ‘look over there’.” (Munroe 2009). Thus, a key step in establishing that the correlation uncovered reflects our biological hypothesis, rather than happenstance, would be to evaluate the generality of this pattern. Some such evidence already exists – similar results have been seen in RFLP based studies of a few R genes in big bluestem and switchgrass (Rouse et al 2011, Zhu et al 2013). Testing for this pattern in other *Silphium* species represents a promising direction, as they would provide evolutionarily independent replication, while covering a similar precipitation gradient, and could use the same RenSeq baits developed here. Extending this study to more distant taxa would provide further evidence supporting this hypothesis.

Functional studies of these R-genes would provide more evidence for our motivating causal hypothesis. A complete, phased, chromosome-level assembly of *S. integrifolium* will both allow for better assessment of whether these genes are functional, and enable association studies to determine the loci of pathogen resistance for incorporation into breeding programs. Additional evaluation of the hypothesis that the number of pathogens affecting Silphium (and/or the variation in their ability to evade a specific R gene), as well as associating specific NLR alleles to resistance to specific pathogens would allow for a more mechanistic understanding of the association uncovered in this paper.

## CONCLUSION

We used RenSeq to sequence the R gene libraries of *S. integrifolium* plants from across a rainfall gradient to evaluate the hypothesis that CMI imposes a biotic stress whose evolutionary pressure should be evident within the genome. We detected a significant positive relationship between CMI and R gene count, supporting our hypothesis. We demonstrate that RenSeq can be used in the evaluation of ecological hypotheses and show promise for the economical interrogation of the specific R gene loci responsible for disease resistance.

## FUNDING

This work was supported by a grant from the National Science Foundations’ Dimensions in Biodiversity Grant [Grant ID: 1738041].

## DATA ACCESSIBILITY AND BENEFIT SHARING

The R gene libraries generated in this study are available on the NCBI SRA database at project accession PRJNA914292.

## AUTHOR CONTRIBUTIONS

KK assembled and annotated the R gene libraries, performed the analyses, generated the figures, and co-wrote the manuscript. KP performed the majority of the field work, common garden experiments, and phenotyping, and co-wrote the manuscript. KT created disease scales and led the recording of disease severity in the field. AH sequenced and assembled the draft genome of *S. integrifolium*. AR designed the RenSeq baits, developed the libraries, and performed the initial steps of bioinformatics. YB, KS, JB, KT and DVT conceived of the project and contributed to the writing.

## ACKNOWLEDGEMENTS

We acknowledge support for this research from NSF Dimension of Biodiversity grant 1738041. This research was supported financially by the Perennial Agriculture Project, a joint project between The Land Institute and The Malone Family Land Preservation Foundation. We gratefully acknowledge financial support from the Foundation for Food and Agriculture Research (FFAR) Fellows program. We thank Eric Cassetta for assisting with all of the field work. We are thankful to Andrew Read and Peter Innes for assistance with code to generate the phylogeny and map figures, respectively.

## COI

Authors declare no conflicts of interest.

## Notes

https://www.ncbi.nlm.nih.gov/bioproject/PRJNA914292

### Competing Interest Statement

The authors have declared no competing interest.

### Summary of Updates

Minor changes to wording in the discussion to reflect changes suggested by reviewers. Primarily, we softened the language regarding the potential generality of the pattern uncovered in this study, as well as propose a simple extension of this natural experiment to cover other cosmopolitan species within the Silphium genus that exist along the same rainfall gradient, and therefore may substantiate the findings herein.

