## Supplementary figures and images for "Disease resistance gene count increases with rainfall in *Silphium integrifolium*"

### FigS1_PacBioTNLsCNLs.png

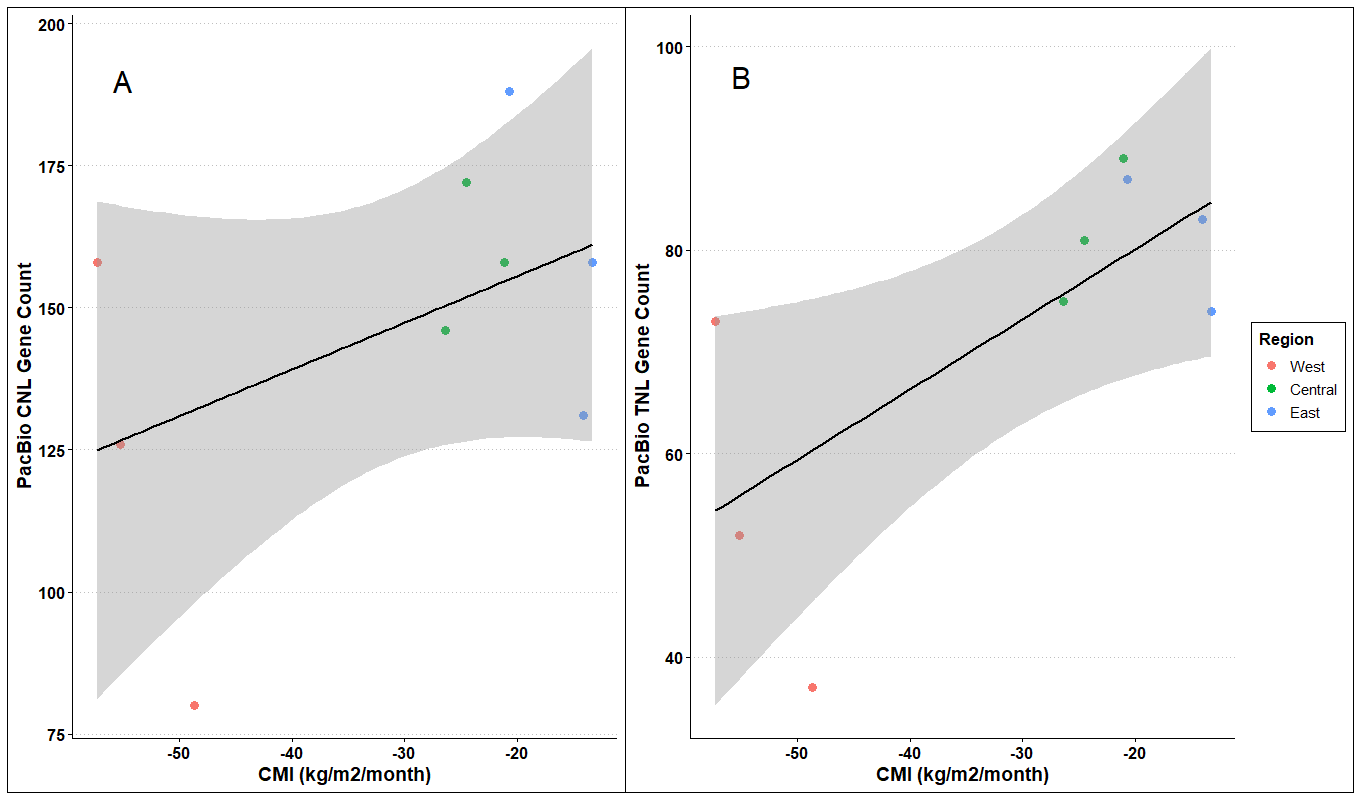

### FigS2_PhenotypesByRegion.png

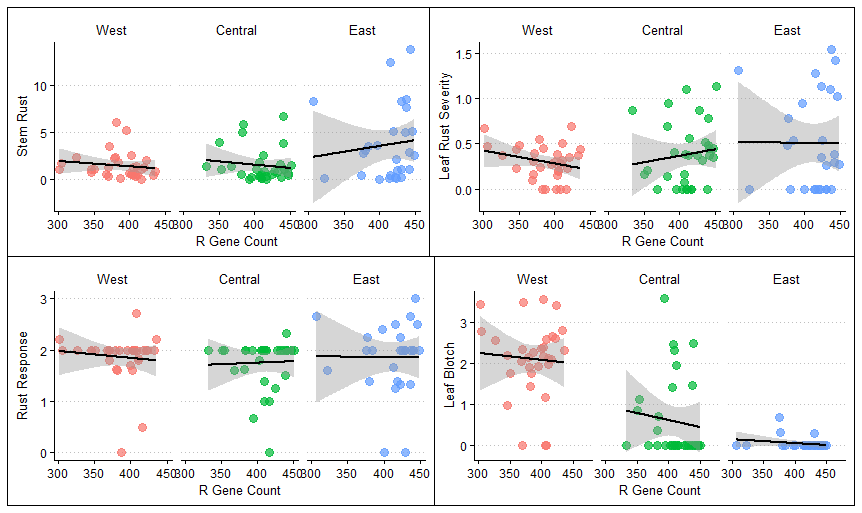

### FigS3_PacBioInputVsCount.png

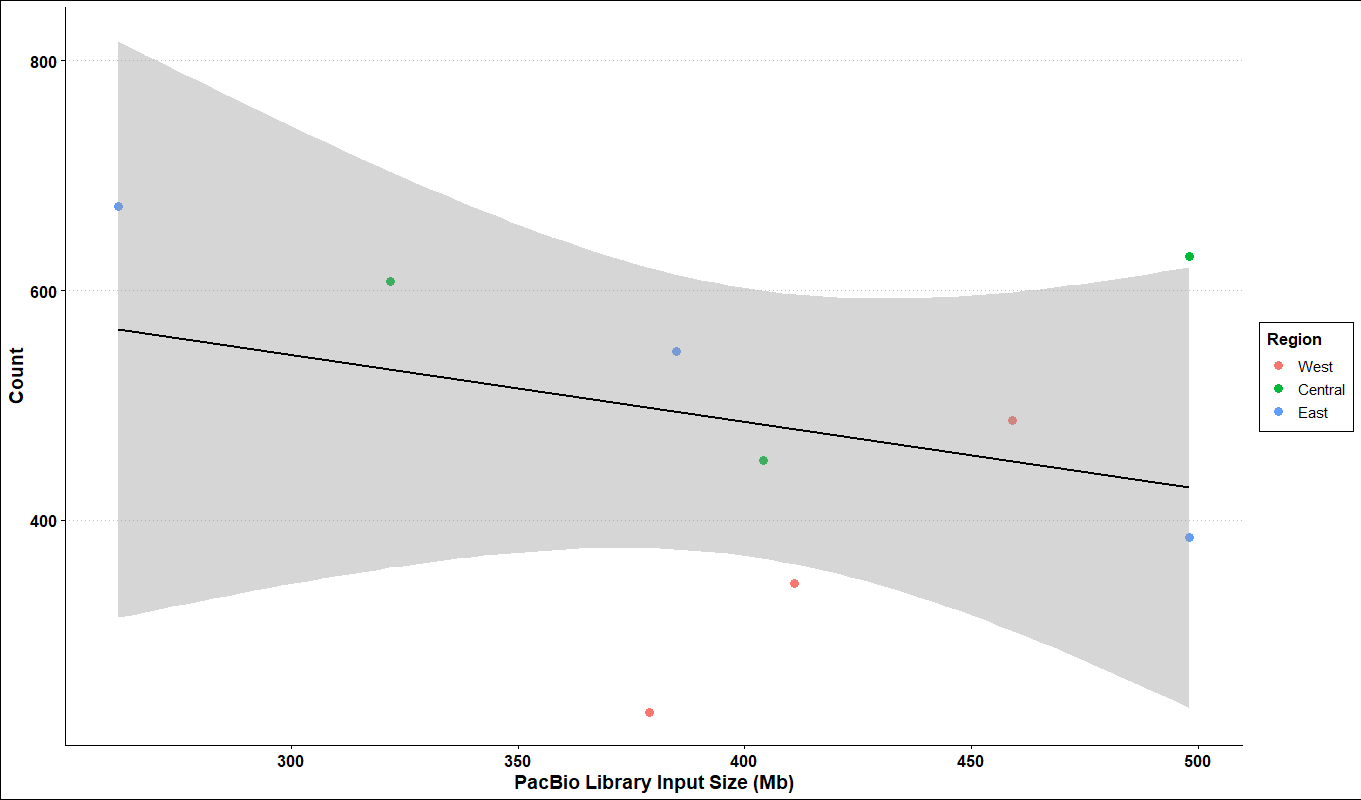

### FigS4_PCoA_Axes.png

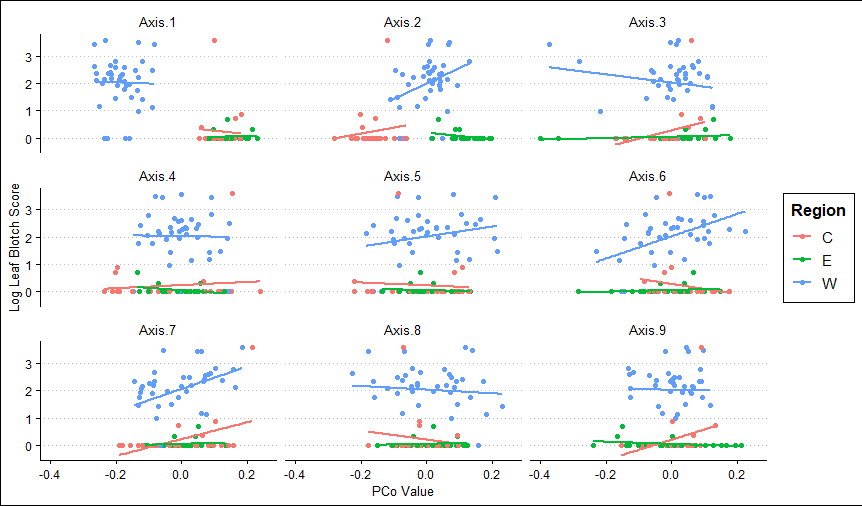
